# CytoSignal Detects Locations and Dynamics of Ligand-Receptor Signaling at Cellular Resolution from Spatial Transcriptomic Data

**DOI:** 10.1101/2024.03.08.584153

**Authors:** Jialin Liu, Hiroaki Manabe, Weizhou Qian, Yichen Wang, Yichen Gu, Angel Ka Yan Chu, Gaurav Gadhvi, Yuxuan Song, Noriaki Ono, Joshua D. Welch

**Affiliations:** Department of Computational Medicine and Bioinformatics, University of Michigan, Ann Arbor, MI, USA; Department of Electrical Engineering and Computer Science, University of Michigan, Ann Arbor, MI, USA; University of Texas Health Science Center at Houston School of Dentistry, Houston, TX, USA

## Abstract

Nearby cells within tissues communicate through ligand-receptor signaling interactions. Emerging spatial transcriptomic technologies provide a tremendous opportunity to systematically detect ligand-receptor signaling, but no method operates at cellular resolution in the spatial context. We developed CytoSignal to infer the locations and dynamics of cell-cell communication at cellular resolution from spatial transcriptomic data. CytoSignal is based on the simple insight that signaling is a protein-protein interaction that occurs at a specific tissue location when ligand and receptor are expressed in close spatial proximity. Our cellular-resolution, spatially-resolved signaling scores allow several novel types of analyses: we identify spatial gradients in signaling strength; separately quantify the locations of contact-dependent and diffusible interactions; and detect signaling-associated differentially expressed genes. Additionally, we can predict the temporal dynamics of a signaling interaction at each spatial location. CytoSignal is compatible with nearly every kind of spatial transcriptomic technology including FISH-based protocols and spot-based protocols without deconvolution. We experimentally validate our results *in situ* by proximity ligation assay, confirming that CytoSignal scores closely match the tissue locations of ligand-receptor protein-protein interactions. Our work addresses the field’s current need for a robust and scalable tool to detect cell-cell signaling interactions and their dynamics at cellular resolution from spatial transcriptomic data.

## Introduction

Intercellular signaling, or cell-cell communication, occurs through many mechanisms, such as when secreted ligands bind to transmembrane receptors or when membrane-bound proteins on adjacent cells dimerize. This communication is crucial for many biological processes in multicellular organisms. For example, cell signaling influences differentiation and fate specification during normal development[1], and intercellular communication coordinates immune response, growth, and physiological tissue function[2] in the developed organism. However, identifying which signaling interactions operate between which cells under particular conditions remains challenging. Single-cell RNA sequencing (scRNA)[3] datasets allowed some progress by revealing which ligands and receptors are expressed by each cell type within a heterogeneous tissue[4]. Numerous methods to infer cell-cell communication from scRNA data have been published, including CellPhoneDB[5], CellChat[6], NicheNet[7], SingleCellSignalR[8], and Scriabin[9], among others. However, these methods have two key limitations: either (1) they do not incorporate information about the spatial proximity of cells; or (2) they infer signaling among groups of cells, rather than at single-cell resolution; or both.

Spatial transcriptomic (ST) protocols measure both gene expression profiles and spatial coordinates within a tissue, providing crucial information about the spatial proximity of cells expressing ligands and receptors. Some computational methods have been developed for detecting cell signaling from ST data, such as CellPhoneDB v4.0[10] and SquidPy[11], but as with most methods for scRNA data, they look for interactions among large predefined groups of cells (cell types).

However, signaling fundamentally reduces to protein-protein interactions between a ligand and a receptor that occur at a particular location in a tissue. The locations at which the interactions occur are in turn determined by the complete spatial microenvironment surrounding the cell; cells do not check the “types” of other cells before interacting. Thus, existing ST methods that detect interactions among cell types are based on abstractions that are necessary for dissociated cell data but limiting when spatial context is available. In short, previous ST computational tools do not make full use of the spatial nature of ST data to infer signaling interactions at cellular resolution. An additional limitation is that existing methods for scRNA and ST data model different types of signaling interactions in the same way. For example, contact-dependent interactions require cells to be immediately adjacent in the tissue[12–14], while interactions between diffusible ligands and transmembrane receptors require cells to be close but not necessarily touching. Furthermore, existing approaches infer only the current state of cell signaling at the moment in time the cells were measured–not how these signaling interactions change over time.

To address these limitations, we developed a novel approach that uses spatially resolved gene expression for estimating static and dynamic signaling among single cells at specific spatial positions. CytoSignal uses a simple, principled score to identify which cells and locations within a tissue have significant activity for a particular signaling interaction. We further developed VeloCytoSignal, a method for predicting the rate of change for a signaling interaction at each tissue location.

Another key challenge in cell-cell communication studies is the validation of inferred signaling activities in real tissues. Previous methods have often relied solely on computational validation, and therefore do not convincingly demonstrate their accuracy and applicability in a real tissue context. In fact, because interactions among cell types are a high-level abstraction, it is difficult to envision what direct validation would look like. In contrast, because CytoSignal predicts something much closer to the mechanism of ligand-receptor interactions, it is possible to validate our predictions more directly. In particular, we designed an *in situ* validation analysis through the proximity ligation assay (PLA) [15–17] to verify the tissue locations at which these interactions occur in the developing mouse embryo. This analysis confirmed the physical locations where a ligand binds to its receptor, providing direct evidence to support CytoSignal’s predictions.

## Results

### CytoSignal and VeloCytoSignal identify locations and dynamics of signaling interactions at cellular resolution

The key idea of CytoSignal is that ligand-receptor signaling is fundamentally driven by a protein-protein interaction that happens at a specific tissue position. This interaction happens, in turn, when all of the requisite ligand and receptor components are expressed in close spatial proximity. Following this assumption, we construct a score to measure the signaling interaction strength at each position within a tissue (**Fig. 1a**). The ligand-receptor interaction score (LRscore) *S* is the product of ligand (*L*) and receptor (*R*) expression in the spatial neighborhood of the cell: *S* = *L* × *R*. Intuitively, this score is motivated by an underlying chemical reaction in which the ligand and receptor proteins interact to form a complex: *L* + *R* → *LR*. Note that the rate of this binding reaction is directly related to the product of ligand and receptor concentration. This strategy gives a simple, interpretable metric of the signaling activity occurring at each position within the tissue.

**Fig.1.**
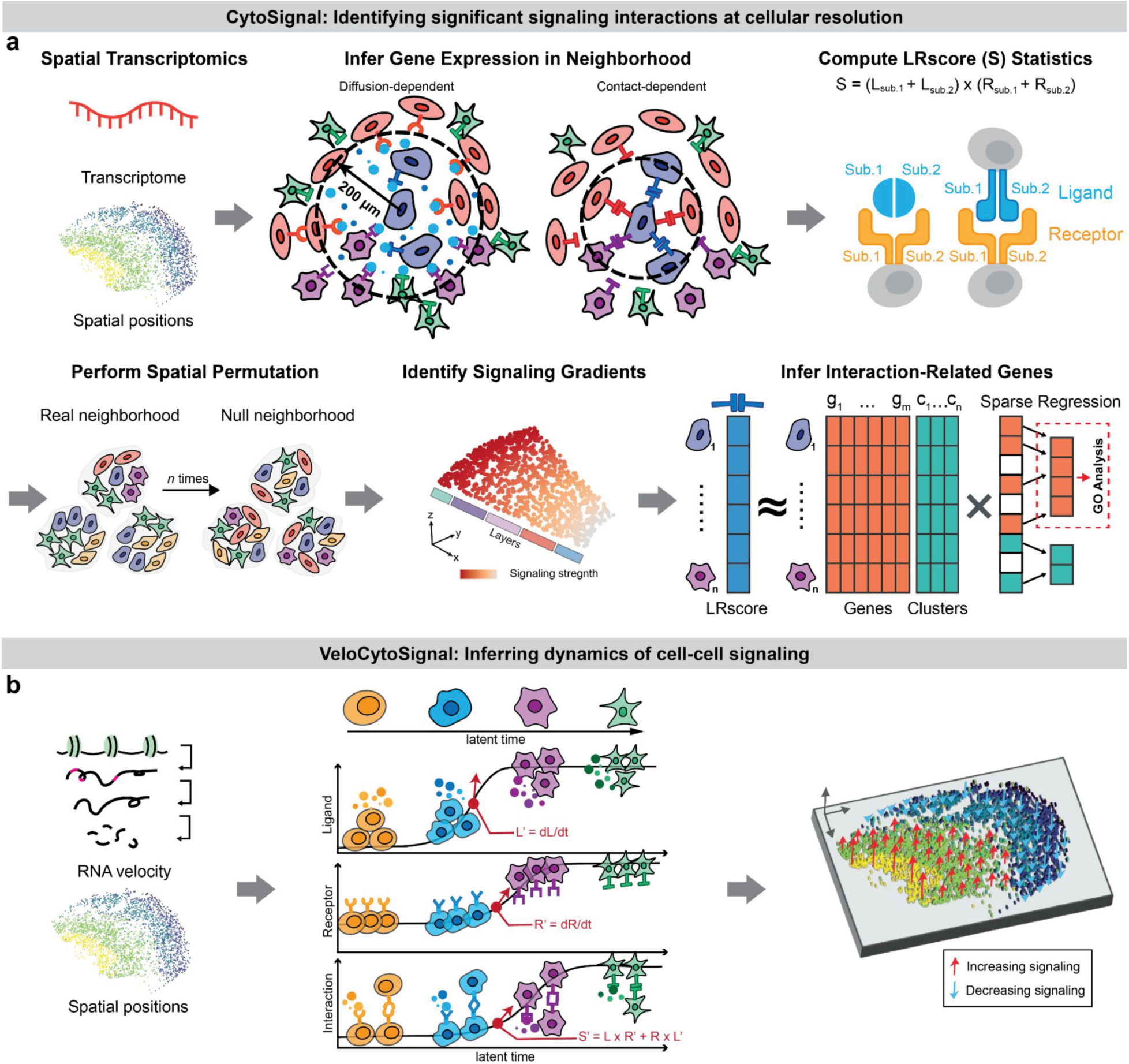
CytoSignal and VeloCytoSignal model the spatiotemporal dynamics of ligand-receptor signaling interactions at cellular resolution. **a**, Diagram of CytoSignal approach. CytoSignal predicts the amount of ligand-receptor protein-protein interaction at each position in a tissue (LRscore *S* = *L* × *R*) for each interaction while considering both the spatial neighbors of each cell and the diffusion- or contact-dependent nature of the interaction. Expression of individual components within interactions involving multiple subunits is also considered. Cytosignal then identifies which cells have significant activity for a particular signaling interaction by spatial permutation. CytoSignal also detects spatial gradients in signaling and signaling-associated differentially expressed genes. **b**, Diagram of VeloCytoSignal approach. VeloCytoSignal predicts the temporal dynamics of a signaling interaction at each spatial location by combining RNA velocity from the ligand and receptor genes to calculate the time derivative (*dS*/*dt*) of the CytoSignal test statistic (*S*). The direction of the arrows indicates whether the signaling activity is increasing (pointing upward, colored in red) or decreasing (pointing downward, colored in blue) in each spatial neighborhood, and the lengths of the arrows indicate the rate of change.

Calculating a signaling score for each spatial position also allows us to separately quantify the locations at which interactions based on diffusible vs. contact-dependent factors occur. We define the spatial neighborhood differently for these two classes of interactions (**Fig. 1a**, Infer gene expression in neighborhood). Contact-dependent interactions require cells to be directly adjacent in the tissue, such as when two or more membrane-bound receptors on different cells dimerize. For contact-dependent interactions, we calculate *L* and *R* for cell *i* using only cells *j* that are immediate spatial neighbors of cell *i*. In contrast, diffusion-dependent interactions can affect cells that are not immediately adjacent to the source cell, though the signal strength still depends on spatial proximity. For diffusion-dependent interactions, we calculate *L* for cell *i* using all other cells *j* weighted by the physical distances between them. In calculating *L*, we use a Gaussian kernel with bandwidth chosen so that most of the kernel density lies within a user-specified radius of the receiving cell (default 200 *μ*m). Next, we use receptor expression to calculate *R*, then calculate *S* using directly connected cell neighbors. In this way, unlike previous approaches, we can quantify the amount of contact-dependent vs. diffusion-dependent signaling happening at each position within a tissue.

Intuitively, our LRscore quantifies the amount of signal received by each cell, which is related to the amount of protein-protein interaction between ligand and receptor occurring at that position. Additionally, because of the simple form of our score, we can infer signal-sending cells for each signal-receiving cell. For contact-dependent interactions, cell *i* is a signal-sending cell for cell *j* and ligand-receptor interaction *k* if *i* expresses the ligand and *j* expresses the receptor. For diffusion-dependent interactions, we determine the signal-sending cell in a similar way, but weight the signal-sending strength by distance (using the same Gaussian kernel described above).

The LRscore serves as a direct estimate for the strength of a signaling interaction at each position in a tissue. However, it is also desirable to know which positions have the most significant evidence for a given signaling interaction. To determine this, CytoSignal constructs a null distribution for *S* by permuting the spatial locations of cells (**Fig. 1a**). We then calculate a one-sided *p*-value for the null hypothesis that the signal strength observed for a particular cell is no larger than expected based on the ligand and receptor expression levels within the tissue. To control for multiple hypothesis testing and potential biases caused by cellular density differences, we further perform spatial false discovery rate correction[18]. Cells with significant signaling activity can then be identified by setting a significance level such as *α* = 0.05. To identify the most interesting interactions, we can rank them by either the number of significant spatial positions or the spatial variability statistic of the LRscore calculated by SPARK-X[19].

CytoSignal can also identify spatial gradients in signaling strength. Because we measure the strength of the signaling interaction at each position in the tissue, independent of discrete cell types, we can readily investigate continuous variation in signaling across a tissue (**Fig. 1a**). Previous approaches that calculate signaling interactions between cell types or using dissociated cell data are not designed to detect this sort of continuous variation.

Another benefit of our approach is that it enables identification of differentially expressed genes associated with a signaling interaction. Because we calculate a score for each signaling interaction at each position in the tissue, we can look for signaling-associated genes in an unbiased fashion. In contrast, previous methods that investigate expression of downstream signaling genes often rely on annotations of signaling pathways[6–8], which can be unreliable and incomplete. To identify differentially expressed genes associated with each signaling interaction, we perform a sparse regression analysis using the output statistic *S* from CytoSignal as the response variable. This also allows us to control for cell type as a confounder when identifying signaling-associated genes. These signaling-associated genes can subsequently be used to identify GO terms or transcription factors associated with the signaling interaction.

We also developed a method of inferring temporal changes in signaling strength at each position in a tissue (**Fig. 1b**). Intuitively, our key insight is that the signaling interaction strength changes when either the ligand or receptor expression changes. Previous work has shown that spliced and unspliced counts can be used to estimate the time derivative of a gene’s expression, a concept called RNA velocity[20–22]. These velocity estimates of the rate of change for ligand and receptor expression can then be used to predict whether a particular signaling interaction will increase or decrease in strength over time. In particular, RNA velocity estimates the time derivative of ligand (*L*) and receptor (*R*) expression individually (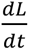 and 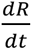). Because the CytoSignal test statistic is simply *S* = *L* × *R*, we can calculate the time derivative of *S* using the product rule of differential calculus:

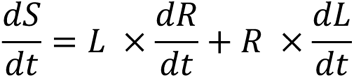

The quantity 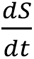 then gives the direction and magnitude of the predicted change in signaling strength for each cell; positive 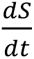 indicates increasing signal strength, while negative 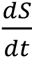 means the signal strength will decrease over time. Of course, this approach relies on having accurate estimates of 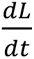 and 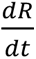 for individual genes. Single-gene velocity estimates from scVelo[21] and velocyto[20] are often unreliable [23], but we utilized our previously published tool, VeloVAE[24], which uses gene-shared latent times to calculate much more accurate velocity estimates for individual genes. We refer to our approach for calculating the rate of change of signaling interactions as VeloCytoSignal, since it merges velocity analysis with signaling analysis.

### CytoSignal detects spatially resolved signaling interactions, spatial gradients, and signaling associated-genes in Slide-seq and Slide-tag data

We first applied CytoSignal to spatial transcriptomic data captured by Slide-seq V2 and Slide-tags from embryonic mouse brains. Slide-seq is a barcoded spatial transcriptomic protocol that measures transcriptome-wide expression of RNAs on 10 μm beads[25]. Slide-tags captures single nuclei after tagging them with spatially mapped barcodes, thus ensuring that transcripts sharing a spatial barcode come from the same cell[26]. Both protocols enable transcriptome-wide detection of RNAs and achieve single-cell (or near single-cell) resolution.

Using these datasets, we performed several analyses that are now possible with cellular-resolution, spatially-resolved signaling inference (**Fig. 2**). In particular, we identified spatial gradients in cell signaling; determined signaling-associated differentially expressed genes; and mapped spatial differences in the amount of contact-dependent vs. diffusion-dependent signaling. We demonstrate these new types of analyses and visualizations using several of the most significant signaling interactions inferred by CytoSignal.

**Fig. 2.**
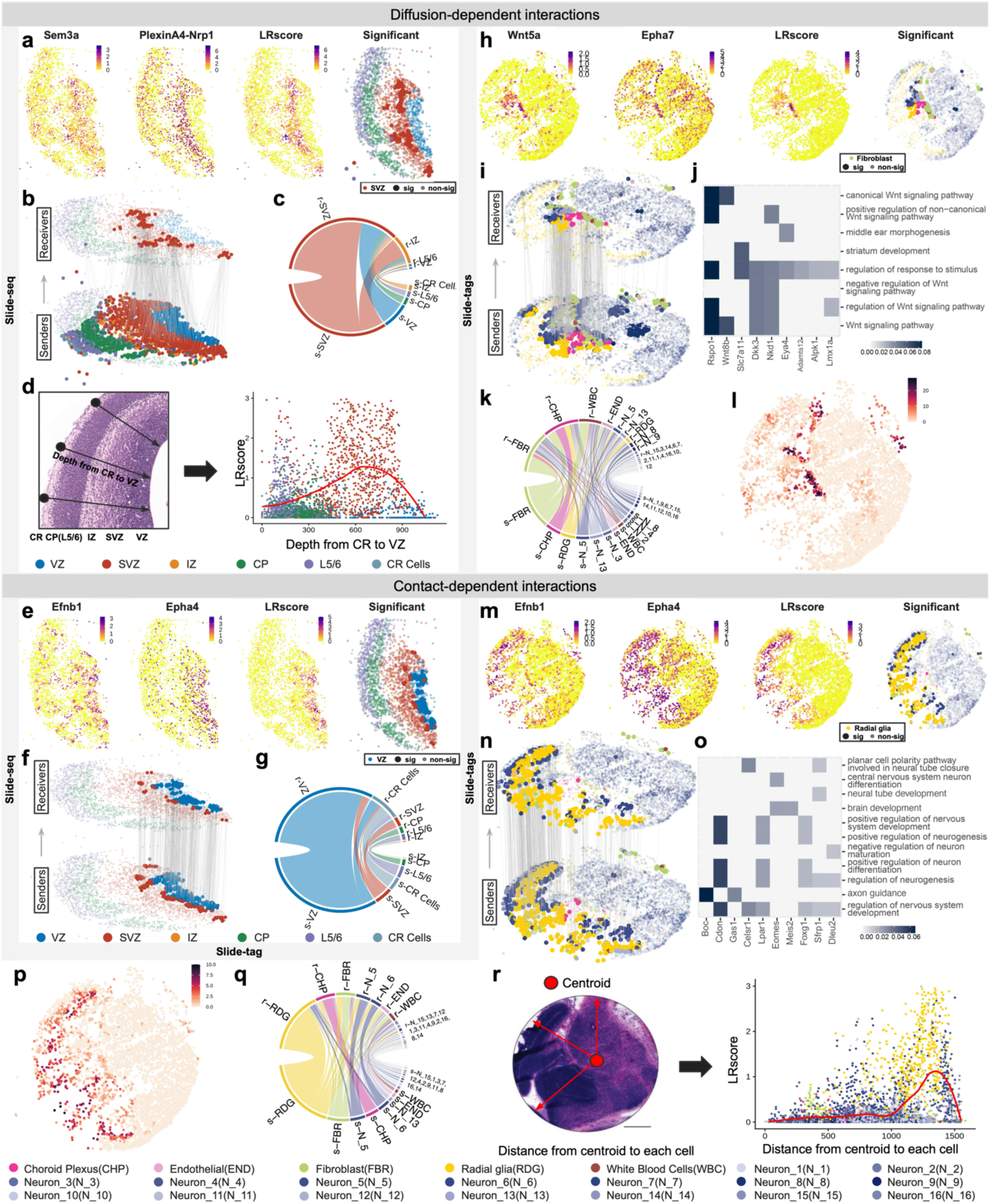
CytoSignal identifies spatial gradients in signaling strength and signaling-associated genes in mouse cortical development. **a,** Ligand (Sem3a) and receptor (PlexinA4 complex1) expression data, Sem3a-PlexA4 signaling interaction scores (LRscores), and inferred locations with significant signaling activity. Each dot is a Slide-seq bead, the x and y coordinates are spatial position, and color indicates expression level, LRscore, or cell type. **b,** Inferred signal-sending and signal-receiving cell interactions for Sem3a-PlexA4. Top layer: locations with significant signaling activity inferred by CytoSignal (signal-receiving cells). Bottom layer: signal-sending cells. **c,** Circos plot showing cluster level summary of signaling activities between tissue locations across all diffusion-dependent interactions. Line thickness indicates the number of significant diffusion-dependent interactions between cells in the corresponding clusters. Clusters with “s-” and “r-” prefixes represent clusters serving as the signal-sending cluster or signal-receiving cluster. **d,** Spatial gradient of LRscores for the Sem3a-PlexinA4 interaction along cortical layers. Each dot represents a bead in the Slide-seq dataset, the x-axis is the cortical depth in µm, and the y-axis is the LRscore. Colors indicate cluster assignments. **e,** Efnb4 (ligand) and Epha4 (receptor) expression, Efnb1-Epha4 signaling interaction scores (LRscores), and inferred locations with significant signaling activity. Each dot is a Slide-seq bead, the x and y coordinates are spatial position, and color indicates expression level, LRscore, or cell type. **f,** Inferred signal-sending and signal-receiving cell interactions for the Efnb1-Epha4 interaction in the Slide-seq cortex data. **g,** Circos plot across all contact-dependent interactions in the Slide-seq cortex data. **h,** Wnt5a and Epha7 expression, Wnt5a-Epha7 LRscores, and inferred locations with significant signaling activity. **i,** Inferred signal-sending and signal-receiving cell interactions for the Wnt5a-Epha7 interaction in the Slide-tags brain data. **j,** Heatmap representation of genes and GO terms associated with the Wnt5a-Epha7 interaction. The color of each grid square represents the regression weight of each gene (positive weight means positive association with Wnt5a-Epha7 signaling). White indicates that this gene is not annotated with the corresponding GO terms in the column. **k,** Circos plot across all diffusion-dependent interactions in the Slide-tags brain data. **l,** Total number of significant diffusion-dependent interactions per spatial location in the Slide-tags brain data. **m,** Efnb1 and Epha7 expression, Efnb1-Epha7 LRscores, and inferred locations with significant signaling activity. **n,** Inferred signal-sending and signal-receiving cell interactions for the Efnb1-Epha7 interaction in the Slide-tags brain data. **o,** Genes associated with the Efnb1-Epha7 interaction and their related GO terms. **p,** Total number of significant contact-dependent interactions per spatial location in the Slide-tags brain data. **q,** Circos plot across all contact-dependent interactions in the Slide-tags brain data. **r,** Efnb1-Epha7 interaction strength as a function of distance from the centroid of the brain.

The output of CytoSignal is a score for each signaling interaction at each individual spatial location. Thus, we can visualize a single signaling interaction using a spatial plot where each point is a tissue position with measured gene expression (bead, spot, or cell), and we color the point by the inferred signaling activity (**Fig. 2a, h, e, and m**). We also developed a novel 3D visualization method (see **Methods**), depicting signal-sending cells (**Fig. 2b, i, f and n**, bottom layer) and signal-receiving cells (**Fig. 2b, i, f and n**, top layer). Each cell is colored by its cluster assignment, and the edges between cells represent inferred signaling interactions. To provide a cluster-level perspective, we utilized Circos plots to show the total number of significant edges sent and received from each cluster across all diffusion-dependent and contact-dependent interactions separately **(Fig. 2c, g, k and q**).

The top interactions inferred by CytoSignal are known to be highly significant in embryonic neural development. In the cerebral cortex captured by Slide-seq, CytoSignal revealed an interaction mediated by a diffusible ligand Sema3a and PlexinA4-Nrp1 co-receptor complex[27], which has a well-described role in neuronal axon guidance[28] and neuronal migration[29](**Fig. 2a**). In the embryonic mouse brain captured by Slide-tags, CytoSignal identified another diffusible-ligand-dependent interaction, Wnt5a-Epha7, that is potentially involved in the patterning decisions in the embryonic cortical areas[30,31](**Fig. 2h**). The contact-dependent interaction between Efnb1 and Epha4 was identified in both datasets and is known to be involved in axon guidance and nerve regeneration[32](**Fig. 2e and m**).

By inferring a signal strength at each spatial position, CytoSignal is able to identify continuous spatial gradients of signal strength. To demonstrate this capability, we calculated the LRscore of the interaction between Sema3a and Plexina4-Nrp1 complex in each bead as a function of its cortical depth (see **Methods**) (**Fig. 2d**). The LRscore of this signaling interaction shows a unique spatial gradient which transiently increases, reaches its highest level in the SVZ layer, and finally decreases. We then performed a similar analysis in the slide-tags data by plotting the LRscore of the interaction Efnb1-Epha4 in each nucleus against its distance from the centroid of the tissue (**Fig. 2r**). This signaling interaction also shows a spatial gradient of LRscores that remains low for neurons near the centroid but increases rapidly with distance from the centroid (**Fig. 2m and 3n**).

**Fig. 3.**
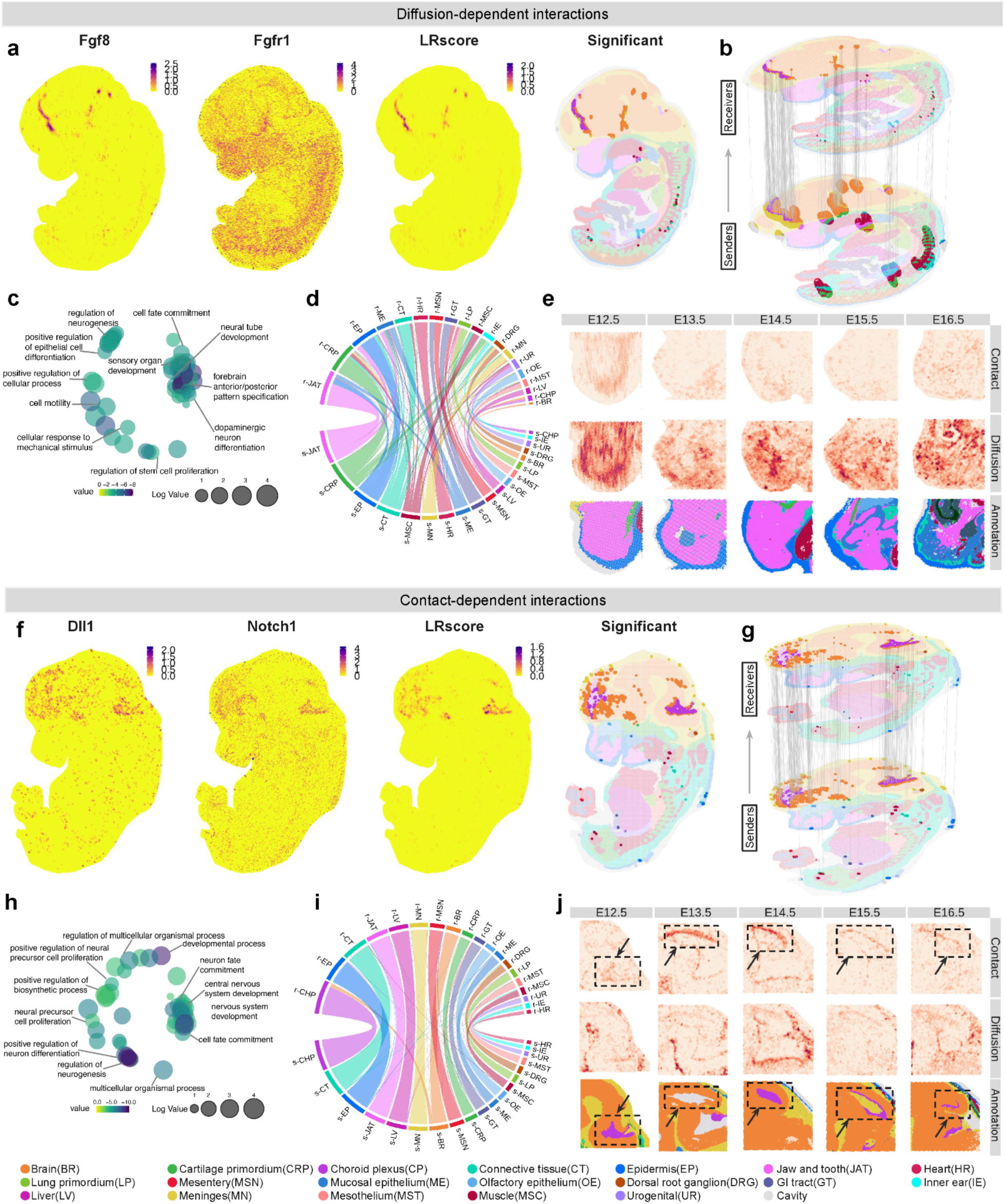
CytoSignal reveals spatiotemporal variation between diffusion and contact-dependent interactions at cellular resolution. **a,** Ligand (Fgf8) and receptor (Fgfr1) expression data, Fgf8-Fgfr1 signaling interaction scores (LRscores), and inferred locations with significant signaling activity. Each dot is a tissue position, the x and y coordinates are spatial positions, and color indicates expression level, LRscore, or cell type. **b,** Inferred signal-sending and signal-receiving cell interactions for the Fgf8-Fgfr1 interaction. **c,** REVIGO plot of GO terms enriched among genes associated with Fgf8-Fgfr1 interaction. **d,** Circos plot of total diffusion-dependent interactions between tissue locations for all clusters. **e,** Total number of significant ligand-receptor interactions per location focusing on the jaw region across five time points from E12.5 to E16.5. Each dot represents a tissue location in each dataset. The first two rows are colored by the total counts of contact-dependent or diffusion-dependent interactions while the third row is colored by cell type annotations. **f,** Ligand (Dll1) and receptor (Notch1) expression data, Dll1-Notch1 signaling interaction scores (LRscores), and inferred locations with significant signaling activity. **g,** Inferred signal-sending and signal-receiving cell interactions for the Dll1-Notch1 interaction. **h,** REVIGO plot of GO terms enriched among genes associated with Dll1-Notch1 interaction. **i,** Circos plot of total contact-dependent interactions between tissue locations for all clusters. **j,** Total number of significant ligand-receptor interactions per location focusing on the brain region across five time points from E12.5 to E16.5. Dotted rectangles highlight the regions that best reflect differences in spatial enrichment between diffusion- and contact-dependent interactions.

We next asked what differentially expressed genes are associated with these signaling interactions. For each interaction, we fit an elastic net regression model to predict the LRscore from gene expression and cluster labels, then select the features with nonzero coefficients that are most predictive for the LRscore (**Fig. 1**). By incorporating cluster labels into the regression model, we can control for cell type as a confounder when identifying signaling-associated gene expression. Note that to avoid circularity we do not include the ligand or receptor genes as predictors in the regression model for a given interaction, since these genes were used to calculate the LRscore, the dependent variable in the regression analysis. We then perform gene ontology (GO) enrichment analysis of the genes with nonzero coefficients. To present these results more clearly, we plot the regression weights of the selected genes as a heatmap along with their enriched GO terms (**Fig. 2j,o**). For the diffusion-dependent interaction Wnt5a-Epha7, interaction-related genes selected by the model are strongly enriched for Wnt signaling terms including “regulation of Wnt signaling pathway” and “canonical Wnt signaling pathway” (**Fig. 2j**). For contact-dependent interaction Efnb1-Epha4, interaction-related genes are strongly enriched for GO terms associated with nervous system development including “positive regulation of neuron differentiation” and “neural tube development” (**Fig. 2o**).

CytoSignal can also identify spatial regions that are enriched for contact-dependent or diffusion-dependent interactions. To demonstrate this, we summarized the total number of significant contact-dependent and diffusible interactions per spatial location (**Fig. 2l and 2p**). Across the tissue, spatial locations near fibroblasts and their adjacent cells seem to have the largest number of significant diffusible interactions (**Fig. 2l**). Circos plots also identify fibroblasts, which are located near choroid plexus, to be both the most active signal-sending and signal-receiving cluster across all diffusible interactions (**Fig. 2k**). This is consistent with biological knowledge that CNS fibroblasts are largely involved in a variety of diffusible signaling pathways including TGFβ, PDGFRβ, and IFNγ signaling[33]. Furthermore, CytoSignal highlights that spatial locations including radial glia beads and their surrounding neurons have the largest number of significant contact-dependent interactions compared with other locations in the tissue (**Fig. 2p**). This aligns with a recent study demonstrating that the contact between radial glia cells and neurons is vital for directing axon-dendrite polarization[34]. Circos plots also provide additional support that radial glia serves as the most intensive signal-sending and signal-receiving cluster (**Fig. 2q**). Another study has shown that radial glial cells constantly interact with each other, which contributes crucially to the formation of the cerebral cortex in the mouse embryonic brain[35].

### CytoSignal reveals locations and signaling-associated genes for contact-dependent and diffusible signaling interactions in mouse embryo Stereo-seq data

We next applied CytoSignal to spatial transcriptomic data measured by Stereo-seq from whole mouse embryos, captured at 8 different time points ranging from embryonic day 9.5 (E9.5) to 16.5 (E16.5)[36]. Stereo-seq uses DNA nanoball-patterned arrays to measure transcriptome-wide expression at sub-cellular resolution. As with the Slide-seq and Slide-Tags data, we used CytoSignal to quantify the strength of signaling interactions at individual tissue locations, which allowed us to identify signaling-associated genes and differences in the locations of diffusible vs. contact-dependent signaling interactions.

Using the Stereo-seq data, CytoSignal was able to detect spatially significant and biologically meaningful interactions. One of the clearest examples was the diffusion-dependent interaction Fgf8-Fgfr1 in the E12.5 E1S1 slice (**Fig. 3a**). Fibroblast growth factor (FGF) signaling mediated by FGF ligands including Fgf8 and their receptors FGF receptors (FGFRs) including Fgfr1 has been well described for patterning and neurogenesis [37–40]. Both LRscore and permutation tests indicated strong activity of the Fgf8-Fgfr1 interaction near the choroid plexus, which is located at the border between the meninges and brain (**Fig. 3a**). Using the LRscores at individual tissue positions, we identified signaling-associated differentially expressed genes. These genes were strongly enriched in GO terms associated with neural development (**Fig. 3c**). Additionally, previous studies have demonstrated the essential role of Fgf8 and Fgfr1 in neural tube development [41,42], which is consistent with the GO term “neural tube development” that is enriched among the signaling-associated genes. The role of Fgf8 in neocortex patterning has also been described previously [43], which is consistent with the GO term “forebrain anterior/posterior pattern specification” that is enriched among the signaling-associated genes.

A clear example of a contact-dependent interaction identified by CytoSignal is Dll1-Notch1 in the E12.5 E1S2 slice (**Fig. 3f**). The Notch-Delta signaling system plays a vital role in the development of the central nervous system [44,45]. For example, sustained expression of Notch1 triggers choroid plexus tumor development in mice [46,47]. This aligns with our results, which indicated an enrichment of Dll1-Notch1 in the right half of the choroid plexus cluster and the intersection area between the Brain cluster and the left half of the Choroid plexus cluster (**Fig. 3f**). Using the LRscores at individual tissue positions, we identified signaling-associated differentially expressed genes. These genes were strongly enriched for GO terms correlated with CNS development including “central nervous system development” and “neuron fate commitment” (**Fig. 3h**). Consistent with these results, it has also been shown that Notch1 in activated forms promotes progenitor cell identity in the mouse embryonic forebrain [48], which corresponds to GO terms including “positive regulation of neuron differentiation” and “neural precursor cell proliferation”.

We also identified interesting patterns in the locations of diffusion-dependent and contact-dependent interactions at the whole embryo scale. For instance, the edge plot for the diffusion-dependent interaction Fgf8-Fgfr1 displays that cells are receiving signals not only from nearby locations but also from more distant locations within adjacent clusters (**Fig. 3b**, bottom layer). As a comparison, the edge plot for contact-dependent interaction Dll1-Notch1 reveals almost straight edges, indicating that cells are primarily communicating with direct neighboring cells (**Fig. 3g**, bottom layer). We also summarized the total number of significant edges sent and received from each cluster across all 209 significant diffusion-dependent and 49 significant contact-dependent interactions separately (**Fig. 3d and Fig. 3i**). We identified the spatial locations near jaw and tooth and epidermis to be the most active signal-sending and signal-receiving region (**Fig. 3d**). Strikingly, diffusible interactions seem to dominate over the contact-dependent ones in the spatial region near the jaw when comparing the total number of significant interactions within each cell across five timepoints (**Fig. 3e**). This aligns with the known regulatory mechanisms of bone development in the jaw and teeth that heavily rely on famous signaling pathways mediated by diffusible molecules, such as Shh, BMP, FGF, and WNT pathways [49–52]. Similarly, from the circos plot, the communication through contact-dependent interactions in the spatial region near choroid plexus and brain exceeds those found in other locations (**Fig. 3i**). Comparing across timepoints, the total number of significant contact-dependent interactions within each cell in this region is nearly at the same level as diffusion-dependent interactions (**Fig. 3j**). However, contact-dependent interactions are primarily concentrated near the boundary between the choroid plexus cluster and the brain cluster. This suggests that contact-dependent interactions have a vital role in guiding the development of border epithelial cells and nearby embryonic neural tissue during development.

### *In situ* proximity ligation confirms locations of ligand-receptor protein-protein interactions predicted by CytoSignal

A key challenge in computational analysis of ligand-receptor signaling is the lack of ground truth to evaluate predictions and the difficulty of direct validation. Previous methods often rely solely on computational validation, which is not sufficient for demonstrating their robustness and applicability. However, the simple, direct nature of the ligand-receptor scores calculated by CytoSignal allows direct validation in real tissues. Specifically, the detection of protein-protein interactions *in situ* can be accomplished using the proximity ligation assay (PLA) (**Fig. 4a**). PLA involves the use of two primary antibodies that specifically bind to the target proteins, followed by oligonucleotide-labeled secondary antibodies (PLA probes). When the two proteins of interest are in close proximity (40 nm), the PLA probes can hybridize with connector oligos and generate a detectable fluorescence signal. Thus, in the context of ligand-receptor interactions, PLA provides direct evidence that both proteins are co-localized within close spatial proximity.

**Fig. 4.**
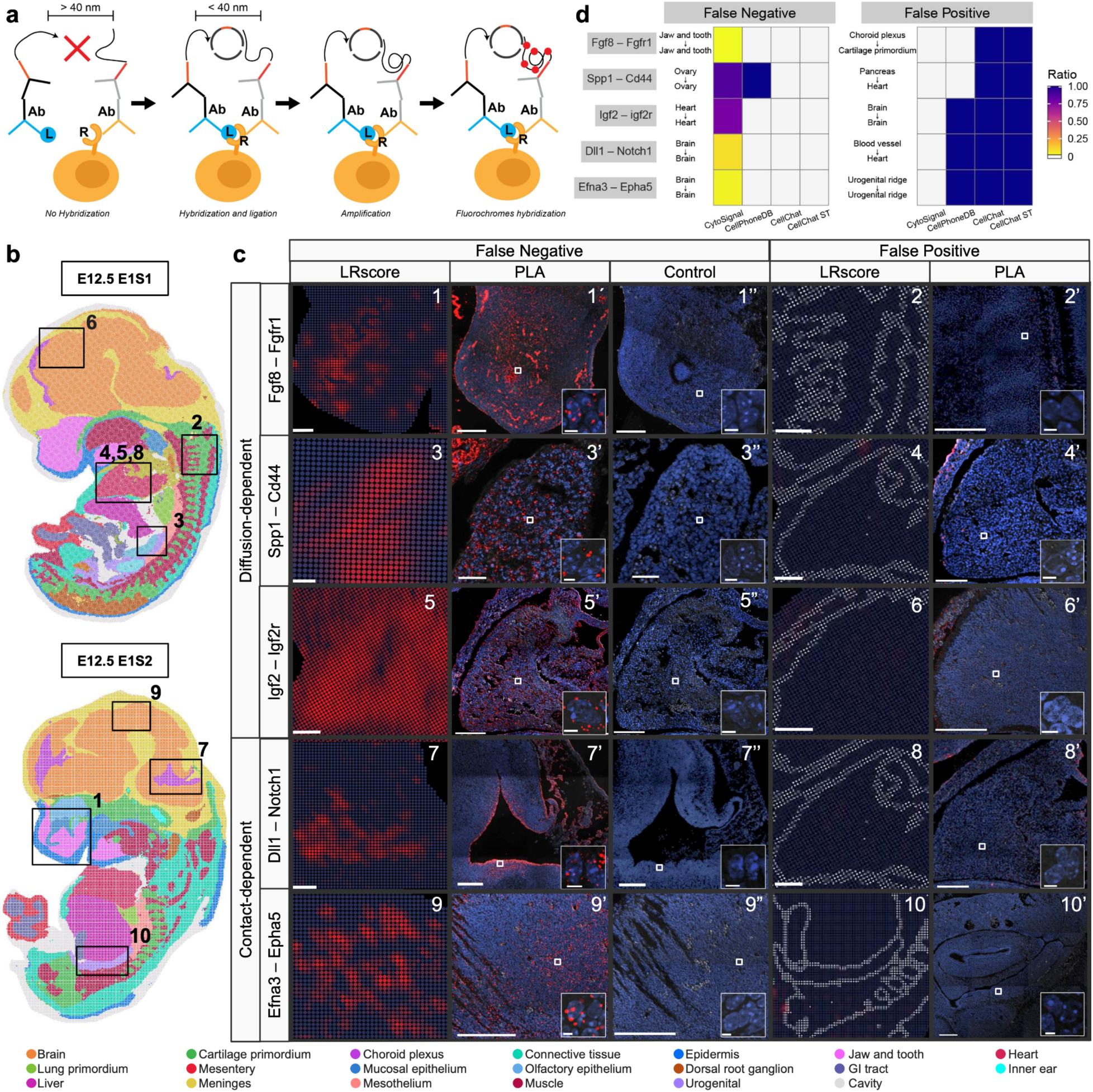
Proximity ligation assay confirms *in situ* ligand-receptor protein-protein interaction locations recovered by CytoSignal. **a,** Diagram of proximity ligation assay (PLA). PLA detects protein-protein interactions using two primary antibodies that specifically bind to the ligand and receptor proteins, followed by PLA probes. The PLA probes bind the primary antibodies and create a fluorescent signal only when the ligand and receptor proteins are within 40 nm of each other. **c,** Fields of view used for PLA imaging and comparison with CytoSignal’s predictions. Numbers colored in black correspond to the numbers colored in white at the upper right corner of **c**. **c,** PLA imaging results and CytoSignal predictions. Each row represents the predictions of CytoSignal and PLA signals for one ligand-receptor interaction. The “LRscore” column shows CytoSignal predictions, and column “PLA” and the “Control” column represents PLA signals in the experimental and negative control samples, respectively. Columns labeled “False Negative” and “False Positive” indicate fields of view where CytoSignal’s predictions are correct but cluster-level methods report false negatives or false positives. The number in white indicates the FOV; numbers correspond to panel **b**. Inset images are high-magnification (63X) views. Scale bar: 200 µm. Scale bar for Ostp-Cd44: 100 µm. Scale bar for inset images: 5 µm. **d,** Quantitative summary of CytoSignal benchmarking with previous methods using PLA validated signaling activities. Each grid square represents the connectivity ratio of each interaction inferred by CytoSignal or previous methods in a pair of clusters. Scores range from 0 (gray) to 1 (blue).

To validate CytoSignal predictions in real tissues, we performed PLA on E12.5 mouse embryos and compared the locations of the PLA signal with the CytoSignal interaction scores predicted from Stereo-Seq data (**Fig. 4a**). We selected five representative interactions with both a large number of significant locations and spatial variation in signaling activity across the embryo. These interactions include three diffusion-dependent interactions (Fgf8-Fgfr1, Ostp-Cd44, Igf2-Igf2r) and two contact-dependent interactions (Efna3-Epha5 and Dll1-Notch1). We assayed these interactions using PLA in a total of eight different spatial fields of view (**Fig. 4b**). CytoSignal’s spatially resolved scores represented by the column “LRscore” colored in red closely match the physical locations of ligand-receptor PLA fluorescence signals. In comparison, we included negative control samples to which we only applied primary antibodies of receptors to block PLA probe hybridization. Importantly, PLA signals from negative control samples shown by the column “Control” confirm the absence of background signal. Furthermore, high magnification imaging (**Fig. 4c**) validates that the PLA signals occur on the cell membrane, consistent with protein-protein interaction between extracellular ligands and membrane-bound receptors. This experiment suggests that spatial positions with high CytoSignal scores indeed correspond with the locations of ligand-receptor interactions *in situ* (**Fig. 4c**).

Additionally, these experiments identified clear examples of how previous cluster-level approaches for cell signaling inference often report false signaling interactions (false positives) or fail to detect signaling interactions that are actually occurring (false negatives) (**Fig. 4c**). To demonstrate this, we ran CellPhoneDB, CellChat and CellChat Spatial (CellChat ST) on the same Stereo-Seq dataset from E12.5 mice. To quantify these cluster-level results, we defined and calculated a connectivity ratio (**r**) for each pair of clusters in all five interactions. The connectivity ratio, ranging from 0 to 1, represents the ratio of statistically significant interacting cells compared to the total number of interacting cells within a pair of interacting clusters. CytoSignal infers at cellular resolution and therefore gives different connectivity ratios according to the real signaling activities of each ligand-receptor interaction. However, previous methods only output cluster-level inferences that indicate all cells in the signal-sending and signal-receiving cluster are interacting, and therefore generate binary connectivity ratios that are either 0 or 1 for all cells in a pair of clusters. For all five interactions we investigated using PLA, CytoSignal successfully detected true positive signaling activity supported by PLA validation (**Fig. 4d**, False Negative, column of CytoSignal), but undetectable by cluster-based methods (**Fig. 4d**, False Negative, other columns in gray). Additionally, CytoSignal correctly predicted the absence of signal (**Fig. 4d**, False Negative, column of CytoSignal in gray) for interactions inferred by previous methods but shown to be false positives by PLA validation (**Fig. 4d**, False Positive, other columns in blue). Specifically, the column of “LRscore” demonstrates that CytoSignal identifies no signal–consistent with the PLA results in the “PLA” column–whereas cluster-based methods reported signaling interactions at those positions (**Fig. 4c**, False Positive).

CytoSignal avoids many of the false positives reported by cluster-level methods since it utilizes spatial gene expression at cellular resolution. To further quantify this advantage, we compared the physical distances between sending and receiving cells identified by CytoSignal and previous methods. Because CytoSignal models signaling activities at the cellular level, we directly calculated the distances between each significant pair of signal-sending and signal-receiving cells. For previous methods that infer at the cluster level, we sampled 1000 cells randomly in the sending and receiving clusters separately, randomly paired them, and calculated the distance between each pair of cells.

Our results demonstrate that CytoSignal only detects significant signaling activities occurring within a 200 μm radius for diffusion-dependent interactions and 40 μm for contact-dependent interactions (**Supplementary Fig. 1a**). However, previous methods such as CellChat and CellPhoneDB that do not consider spatial information predict both diffusion-dependent and contact-dependent interactions even in cells over 8000 μm apart (**Supplementary Fig. 1c, 1d**). Importantly, even cluster-level methods that incorporate spatial information such as CellChat Spatial still predict signaling interactions (even contact-dependent interactions) between extremely distant cells (**Supplementary Fig. 1b**). These results clearly demonstrate CytoSignal’s unique advantage of performing cellular resolution signaling inference in the spatial context.

We also benchmarked the time and memory usage of CytoSignal against previous methods on 8 whole mouse embryo datasets with the exact same genes measured by stereo-seq. Despite operating at cellular resolution, CytoSignal scales to large numbers of cells and shows similar time and memory performance with previous cluster-level methods (**Supplementary Fig. 2**). For example, when applied on a dataset of 121,715 cells using a single core, CytoSignal finished running in 6 minutes and used 8,152 MiB of memory.

### CytoSignal detects ligand-receptor signaling activity from 10X Visium, Seq-Scope, MERFISH, and STARmap PLUS data

An exciting advantage of CytoSignal is its wide applicability to many kinds of spatial transcriptomic technology, as long as the inputs are gene expression counts in cell-by-gene matrix format and spatial coordinates in cell-by-spatial format. We applied CytoSignal to a whole-embryo dataset measured by 10X Visium, a protocol that captures transcriptome-wide expression within 55*μ*m spots. Unlike data types we analyzed in previous sections, the spots in 10X Visium usually contain 1-10 cells depending on tissue type (**Fig. 5f**). Nevertheless, CytoSignal is able to quantify signaling at spot level without requiring deconvolution. For diffusion-dependent interactions, since the spot size is smaller than the preset diffusion radius, we can still infer signaling activities for spatial neighbors within a 200 μm radius (**Fig. 5f**). For contact-dependent interactions, because cells will only touch their directly connected cells within the same spot, we only need to consider gene expression within each spot without applying spatial smoothing across spots (**Fig. 5f**).

**Fig. 5.**
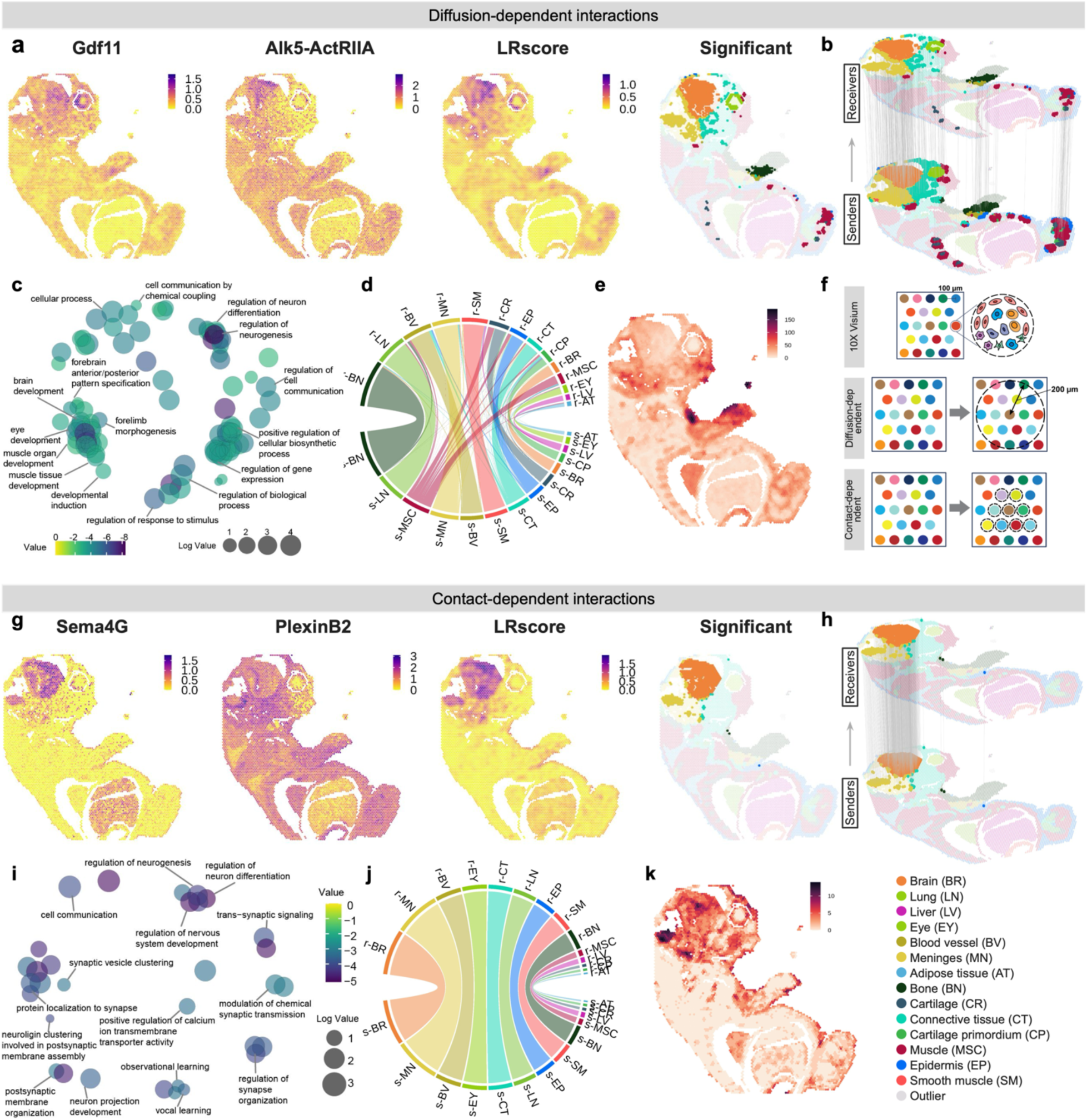
CytoSignal identifies spot-level signaling activity without requiring deconvolution. **a,** Ligand (Gdf11) and receptor (Alk5-ActRIIA complex) expression data, Gdf11-Alk/Act signaling interaction scores (LRscores), and inferred locations with significant signaling activity. Each dot is a tissue position, the x and y coordinates are spatial positions, and color indicates expression level, LRscore, or cell type. **b,** Inferred signal-sending and signal-receiving cell interactions for the Gdf11-Alk/Act interaction. **c,** REVIGO plot of GO terms enriched among genes associated with the Gdf11-Alk5/ActRIIA interaction. **d,** Circos plot of total diffusion-dependent interactions between tissue locations for all clusters. **e,** Total number of significant diffusion-dependent interactions per spot. **f,** Diagram of how LRscores are calculated for Visium spots containing multiple cells. LRscores for contact-dependent interactions are calculated using genes co-expressed within the same spot, while diffusion-dependent interactions are calculated as with cellular-resolution data types using nearby spots weighted by distance. **g,** Ligand (Sema4G) and receptor (PlexinB2) expression data, Sema4G-PlexinB2 signaling interaction scores (LRscores), and inferred locations with significant signaling activity. Ligand (Sema4G) and receptor (PlexinB2) expression data used by CytoSignal and its inferred locations with significant signaling activity. **h,** Inferred signal-sending and signal-receiving cell interactions for the Sema4G-PlexinB2 interaction. **i,** REVIGO plot of terms enriched among genes associated with the Sema4G-PlexinB2 interaction. **j,** Circos plot of total contact-dependent interactions between tissue locations for all clusters. **k,** Total number of significant contact-dependent interactions per spot.

CytoSignal identifies clear examples of spatially constrained ligand-receptor interactions involving diffusible and contact-dependent ligands from 10X Visium data. One example is a well-described diffusion-dependent interaction between the Gdf11 ligand and the Alk5-ActRIIA receptor complex. TGF-β family member Gdf11, activin type I including Alk4, Alk5 and Alk7, and activin type II including ActRIIA/ACVR2A and ActRIIB/ACVR2B together activate the canonical BMP/TGF-*β* signaling pathway and therefore uniquely contribute to CNS development [53][54,55]. Similarly, our results indicate a strong enrichment of both LRscore and significant spots near the brain and meninges (**Fig. 5a, 5b**). Using the LRscores at individual tissue positions, we identified signaling-associated differentially expressed genes. These genes were strongly enriched for GO terms including “brain development” and “forebrain anterior/posterior pattern specification" (**Fig. 5c**). Moreover, Gdf11 negatively controls the number of neurons in the retina [56], and regulates hindlimb development [57] and bone mass [58]. Consistent with this, the interaction is enriched in the developing eye, forelimb, and hindlimb bone, with GO terms including “forelimb morphogenesis” and “eye development”.

CytoSignal also highlights numerous contact-dependent interactions, such as Sema4G-PlexinB2. Class 4 Semaphorins and Plexins are cognate ligand-receptor families that play crucial roles during the development of the mammalian nervous system [59–61]. Previous studies have also described Semaphorin 4G and 4C as ligands of receptor Plexin B2, and these interactions are essential for the migration of cerebellar granule cells in mice [62]. Correspondingly, our results suggest a strong enrichment of both LRscore and significant spots within the brain and part of the meninges (**Fig. 5g and 5h**), with correlated GO terms including “regulation of nervous system development” and “regulation of neuron differentiation” (**Fig. 5i**). Heatmaps of total significant interactions per spot indicate that the locations close to the forelimb and those around the brain and meninges have the largest number of significant diffusible and contact-dependent interactions, respectively (**Fig. 5e and 5k**). Similarly, forelimb bone cells, and brain cells are the most active signal-sending and signal-receiving cell types, respectively (**Fig. 5d and 5j**).

In addition to sequencing-based ST protocols, we have also applied our method to data measured by fluorescence-based protocols including MERFISH [63]. MERFISH is a ST protocol that can simultaneously detect transcripts and their spatial positions at single-molecule resolution. We utilized QuickNii [64] to register the MERFISH data to the Allen Brain Atlas common coordinate framework [65,66] and assign each cell to a brain region. CytoSignal identified multiple significant signaling interactions, including an interaction between the diffusible ligand Sdf1 and the receptor Cxcr4 [67] in the lateral ventricle. Neural progenitor cells (NPCs) located in the subventricular zone (SVZ) play a crucial role in adult neurogenesis [68].

Stromal cell-derived factor Sdf1 is vital for regulating and maintaining both adult and embryonic NPCs [69,70]. Previous studies have shown that the Sdf1-Cxcr4 signaling interaction plays a vital role in adult SVZ cell differentiation and proliferation [71,72]. Correspondingly, in both slices, CytoSignal identified that cells with significant signaling activities are mostly enriched in the lateral ventricle (**Supplementary Fig. 3**).

We then applied CytoSignal to another probe-based protocol called STARmap PLUS [73]. We used a dataset from a mouse model of Alzheimer’s disease (AD). CytoSignal identified multiple significant signaling interactions from among the ligand and receptor genes included in the STARmap PLUS gene panel. One example is an interaction between the diffusible ligand Semaphorin 3a (Sema3a) and its receptor Plexin D1 (Plxnd1) [74]. Semaphorins such as Sema3a are well described to be relevant to neurodegenerative diseases including AD [75]. Our results showed cells with significant signaling activity are mostly enriched near the Cornu Ammonis 1 (CA1), with another minor enrichment in the Dentate gyrus (DG) (**Supplementary Fig. 4a**).

CytoSignal also identified a contact-dependent interaction between Amyloid precursor protein (App) and the transmembrane receptor Cd74. AD is characterized by the presence of extracellular plaques, which are primarily composed of amyloid-β (Aβ) peptides. Previous studies have shown that the interaction of Cd74 and App suppresses Aβ production [76]. Correspondingly, CytoSignal inferred that this interaction has the largest number of cells with significant signaling activities compared to all other interactions, with a nearly ubiquitous distribution of this interaction across spatial locations (**Supplementary Fig. 4b**).

### VeloCytoSignal identifies spatiotemporal dynamics of signaling activity during mouse embryonic development

Cell-cell communication plays a central role in dynamic processes such as cell differentiation. Thus, elucidating spatiotemporal changes in cell signaling activity is crucial for understanding the role of cell-cell communication in cell fate specification and tissue homeostasis. However, no existing computational approaches directly address the question of how to detect temporal changes in signaling from snapshot single-timepoint data. Previous cluster-level methods [21,77,78] based on scRNA-seq data do not model spatial positions and cannot infer the rate of change of signaling over time. Approaches that consider spatial positions only infer at cluster level and also do not model signaling dynamics.

To address these limitations, we developed an approach for detecting spatiotemporal changes in signaling activity that we call VeloCytoSignal. Importantly, our approach can predict dynamics using spatial data from only a single time point. VeloCytoSignal predicts the instantaneous rate of change for a signaling interaction at a specific position within a tissue by combining RNA velocity from the ligand and receptor genes to calculate the time derivative (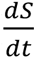) of the CytoSignal LRscore (*S*). One output of VeloCytoSignal is a 3D velocity plot, with the directions of the arrows indicating whether the signaling activity is currently increasing (pointing upward) or decreasing (pointing downward) at each spatial position (**Fig. 6a, 6c, 6e**).

**Fig. 6.**
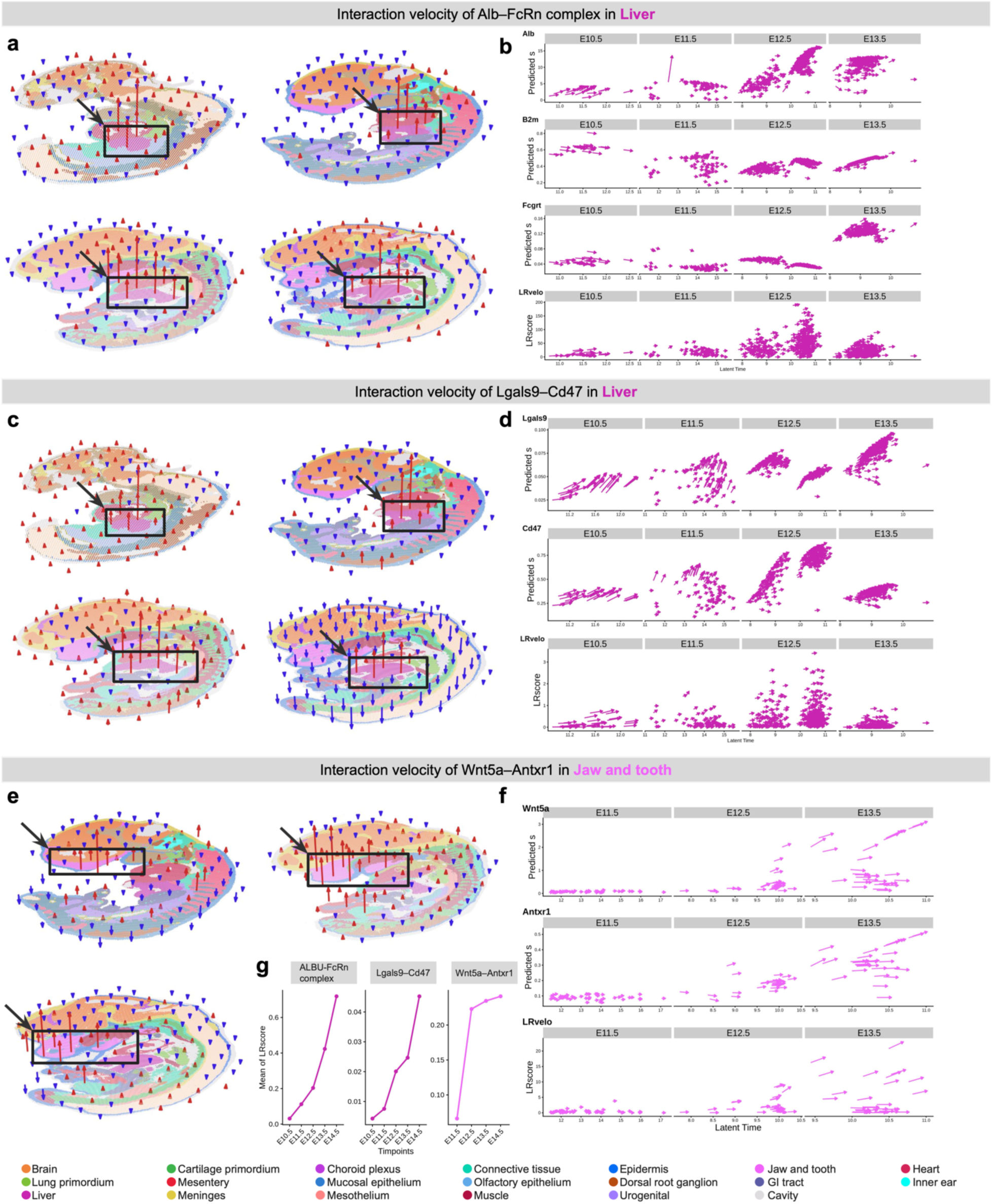
VeloCytoSignal revolves spatiotemporal upregulating signaling activities across multiple embryonic stages. **a,** Signaling velocity (LRvelo) scores for the Alb-FcRn interaction across four time points (E10.5-E13.5). Each point is a spot and color indicates cell type. 3D arrows overlaid on the spatial coordinates indicate the signaling velocity for a spatial bin; arrow direction and color indicate the sign of the velocity (red and up indicate increasing signaling; blue and down indicate decreasing signaling). Arrow length indicates velocity magnitude. Black rectangles highlight the liver. **b,** RNA velocity estimates for ligand Alb and receptor FcRn (calculated by VeloVAE) used as input to VeloCytoSignal and corresponding interaction velocity estimates (LRvelo). Each dot is a stereo-seq position, the x-axis is the inferred latent time, and the y-axis is the spliced expression level for RNA velocity and LRscore for interaction velocity. Arrow direction and length indicate the velocity of each cell. Note that cells are subsampled to avoid overplotting. **c,** LRvelo scores for the Lgals9-Cd47 interaction plotted as in **a**. **d,** RNA velocity estimates for ligand Lgals9 and receptor Cd47 (calculated by VeloVAE) and interaction velocity LRvelo. Black rectangles highlight the liver region. **e,** LRvelo scores for the Wnt5a-Antxr1 interaction plotted as in **a** and **c. f,** RNA velocity estimates for corresponding ligand gene Wnt5a and receptor gene Antxr1 and interaction velocity LRvelo. Black rectangles highlight the jaw and teeth region. **g,** Mean of spatially resolved scores calculated by CytoSignal as a function of time. Dots and lines are colored by cell types.

To validate the predicted temporal trends in signaling activity, we performed VeloCytoSignal on Stereo-Seq data from four consecutive time points (E10.5, E11.5, E12.5, and E13.5). We used each time point separately to predict the temporal trend in signaling activity, thus blinding VeloCytoSignal to the true time point information, then compared the predicted trend with the observed signaling activity at the next time point. To better show the coherence between gene and interaction velocity, we also employed an arrow plot showing the velocity estimates of genes and interactions against gene-shared latent time in downsampled spots (**Fig. 6b, 6d, 6f**). Gene velocity, predicted spliced counts, and latent time in each spot were inferred by our previously published method VeloVAE, and the interaction velocity values (LRvelo) were calculated by VeloCytoSignal.

VeloCytoSignal is able to accurately predict temporal changes in signaling interactions vital for embryonic development. We first investigated a classical diffusion-dependent interaction between albumin (Alb) and neonatal crystallizable fragment receptor complex (FcRn), which is enriched in the liver across multiple timepoints from E10.5 to E13.5. Albumin is a circulating protein produced by hepatocytes of the liver that acts as a multi-functional transporter of essential substances and waste products [79]. FcRn serves as the receptor of Alb and regulates its homeostasis [80,81]. Recent research has revealed that the interaction between Alb and FcRn plays a crucial role in regulating liver homeostasis and response to injury [82]. Accordingly, our 3D velocity plots across four time points showed a sharp increasing trend in Alb-FcRn interaction uniquely in the liver (**Fig. 6a**). Velocity estimates for the ligand Alb along with the receptor components B2m and Fcgrt displayed a consistent trend, with the rate of increase becoming higher at each time point (**Fig. 6b**). We validated that this predicted temporal trend is correct by plotting the average LRscore of the real data across consecutive embryonic time points, which confirmed the trend predicted by VeloCytoSignal (**Fig. 6g**).

Another striking example of signaling dynamics involves an interaction between the diffusible ligand Galectin-9 (Lgals9) and the Cd47 receptor [83]. As with the Alb-FcRn interaction, the VeloCytoSignal prediction of the temporal trend of Lgals9-Cd47 signaling closely matches the true temporal trend observed across the 4 consecutive time points (**Fig. 6c and 6g**). This trend is also consistent with previous findings: Galectin-9 (Gal-9) is a member of the galectin family and particularly enriched in the liver [84]. Additionally, Gal-9 is directly related to the accumulation of macrophages in the liver and reduces hepatocellular damage [85,86]. Cd47 is a transmembrane receptor located on the cell surface that regulates phagocytosis through macrophages in the liver [87,88]. Importantly, VeloCytoSignal is able to leverage the velocity differences between ligand and receptor genes when calculating the velocity of the signaling interaction. For example, Cd47 has a lower but still increasing velocity in E13.5 compared to E12.5, while the velocity of ligand Lgals9 in later time points is consistently higher than previous time points (**Fig. 6d**). These subtle differences are reflected in the 3D velocity plot by shorter arrows and in the arrow plot by lower LRvelo scores.

We also found a signaling interaction with a clear temporal trend in the developing jaw region of the mouse embryo. VeloCytoSignal identified a significant interaction between the diffusible ligand Wnt5a and Anthrax toxin receptor 1 (Antxr1) [83] in the anatomical region that gives rise to the jaw and teeth. Tooth development is a complicated process that includes multiple stages [89]. Several signaling pathways including the WNT pathway are activated at all stages of tooth development [90]. Wnt5a is a representative member of the non-canonical WNT pathway and plays a well-established role in embryonic tooth development [91,92]. Meanwhile, Antrx1 encodes a transmembrane protein and modulates Wnt signaling during vascular development [93], whereas recessive mutations of this gene are closely associated with tooth agenesis[94]. Our VeloCytoSignal analysis predicts that the signaling activity of the Wnt5a-Antxr1 interaction is largely static across the embryo, except for the jaw and tooth region, which shows significant signaling velocity (**Fig. 6e**). This trend is supported by the velocities of both ligand and receptor genes (**Fig. 6f**). Importantly, this trend matches closely with what is observed in the real data from consecutive time points (**Fig. 6g**). Interestingly, the signaling activity of this interaction dramatically increases from E11.5 to E12.5 (**Fig. 6g**). This timing aligns with the known stages of tooth development, in which teeth first become morphologically distinguishable at E11.5, while the cells of the dental lamina proliferate at E12.5, leading to a dramatic increase in WNT signaling activity [95].

## Discussion

Cell-cell communication mediated by ligand-receptor interactions plays a crucial role in cell differentiation, tissue homeostasis, immune response, and disease, but has been difficult to study in an unbiased fashion. CytoSignal infers activities and dynamics of signaling interactions at each spatial position within a tissue using spatial transcriptomic data. By taking this cell-level approach, we can perform several novel types of analyses: identifying spatial gradients in signaling; finding signaling-associated genes; and locating contact-dependent vs. diffusion-dependent interactions. Our model accurately identified interactions in the developing mouse embryo, neurogenesis in the adult mouse brain, and a mouse model of neurodegeneration. Additionally, we confirmed that the interaction scores predicted by CytoSignal correspond to the positions of ligand-receptor protein-protein interaction *in situ*.

CytoSignal and VeloCytoSignal are applicable to most spatial transcriptomic techniques, requiring only a cell-by-gene matrix and a cell-by-spatial-position matrix. We hope that our methods can help solve the current need for analyzing rapidly increasing spatial transcriptomic datasets. As technology for *in situ* molecular measurement continues to advance, we aim to continue expanding applicability of these approaches across new data modalities.

We anticipate that our approach can be used to unveil the signaling networks across a variety of physiological and pathological contexts, in addition to the developmental contexts we considered here. For instance, in the tumor microenvironment, CytoSignal could reveal interactions among malignant cells, healthy cells, and immune cells. Additionally, VeloCytoSignal could provide unique insights into how the dynamics of spatially resolved signaling pathways affect tissue repair over time when applied to time-series data from damaged tissue. We hope that our methods enable similar analyses across a range of tissue contexts.

## Data Availability

Slide-seqV2 data, Slide-tags data and STARmap PLUS data are available at Broad Institute’s single-cell repository (https://singlecell.broadinstitute.org/single_cell/) with ID SCP815, SCP2170 and SCP1375. Stereo-seq datasets from all timepoints are available at https://db.cngb.org/stomics/mosta. The 10X VISIUM dataset is available at https://www.10xgenomics.com/datasets/visium-cytassist-mouse-embryo-11-mm-capture-area-ffpe-2-standard. The MERFISH dataset is available at https://info.vizgen.com/mouse-brain-data.

## Code Availability

CytoSignal and VeloCytoSignal are implemented in R. The package is available on GitHub at https://github.com/welch-lab/CytoSignal.

## Acknowledgements

We gratefully acknowledge Nianxin Yang for discussions on proximity ligation experiments and Fan Feng for insights on Gaussian kernel inference. We also gratefully acknowledge grant support: Chan-Zuckerberg Initiative grant DI2-0000000090 to J.D.W. and J.L. and a Rackham Fellowship to J.L.

## Methods

### Spatially resolved scores of signaling interactions (LRscore)

The spatially resolved interaction score of each interaction in each location is defined as the co-expression of ligand and receptor genes within close spatial proximity. The computation can be divided into three main steps: 1) defining spatial neighbors for each location; 2) calculating the amount of ligand (*L*) and receptor (*R*) each location can receive from their spatial neighbors; and 3) calculating ligand-receptor co-expression within each spatial neighborhood.

CytoSignal defines spatial neighborhoods for diffusion-dependent and contact-dependent interactions differently. For diffusion-dependent interactions, for each location *i*, we define its spatial neighbors as all locations *j* within a circle centered on location *i* with a predefined radius *r* (200 µm by default). For the amount of receptors expressed at each location (*R*), we directly use the normalized gene expression without smoothing since receptors are located on cell membranes and are not diffusible. We next calculate the amount of ligand *L* that location *i* receives using all other locations *j* weighted by the physical distances between them transformed by a Gaussian kernel. For determining the parameters of the kernel, we assume that a ligand diffuses across a two-dimensional tissue space and its concentration follows a Gaussian distribution, with its overall concentration summing up to unity. We want to determine a standard deviation σ such that in the space beyond a predefined radius *r*, the total concentration is less than a trivial value ε. In other words:

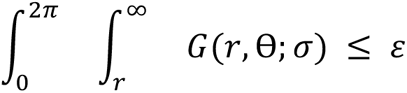

The solution to the above inequality is the following:

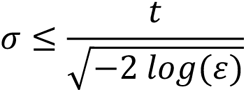

For predefined *r* = 200 µm and ε = 0.001, we can get that *σ_max_* = 53.8 µ*m*.

For contact-dependent interactions, we utilize Delaunay Triangulation (DT) to find the directly connected neighbors of each cell. To exclude outliers, we also filtered out DT neighbors that are too distant from the center cell, using a default threshold of 200 µm. For calculating the amount of ligands received at each location, we calculate a weighted average of gene expression across the neighborhood, including the index location itself. The overall weight sums up to 2, with the weights of the DT neighbors adding up to unity and the weight of index location also being unity. Finally, for calculating the LRscore of each interaction within each spatial neighborhood, we multiply *L* and *R* within each location and apply an average within its DT neighborhood.

### Regression analysis and gene enrichment analysis

For Stereo-seq data, we first subset the genes to 1239 TFs using a 1375 TFs list. An elastic net with identity link function is then fitted using the “*glmnet*” package in R with all selected TFs as well as all cluster labels. We tuned the model with 6 different mixing parameters (0.5, 0.6, 0.7, 0.8, 0.9, 1) by cross validation. The best model is used to choose the most interaction score predictive TFs for each interaction.

For Slide-tag data and Visium data, we filter genes expressing in less than 50 cells as well as the ligand and receptor genes. We iteratively apply one-sided wilcoxon-tests between inferred significant and insignificant cells within every cluster, with an alternative hypothesis that the significant cells have higher expression levels. For each cluster, top 50 significant genes (Benjamini–Hochberg correction applied, FDR < 0.05) from the Wilcoxon-test are selected as the predictors for the downstream analysis. For high-quality data with lower sparsity, we utilize all available transcription factors (TFs) from a publicly available TF list[96] as predictors. We do not apply any filtering steps on the TFs since they are usually well-described and relatively few in number. Cluster annotations are also considered as covariates and are added to all regression models.

The output of regression is a set of predictive TFs or genes as well as their coefficients which are then passed to downstream analyses such as gene ontology (GO) enrichment analysis. The GO analysis by GOrilla can be done by the following settings: (1) organism: “Mus musculus”; (2) running mode: “Two unranked lists of genes”, with model selected TFs as the target set and the other TFs in the 1239 TFs be the background for the Stereo-Seq. For Slide-tag and Visium, we used model selected genes as the target set and other genes in the dataset as the background. (3) ontology: “Process”; (4) check the box “Show output also in REVIGO”. For the “P value threshold”, we used 0.001.

For visualizing the signaling associated genes and their enriched GO terms by heatmap, we manually pick significant GO terms (p-value = 0.001) according to their functions. All elastic net selected genes are ranked by the regression weights and at most top 20 genes are shown in the heatmap.

### Calculating signaling interaction velocity (LRvelo)

In VeloCytoSignal, the velocity of ligand-receptor interaction requires RNA velocity, which is based on mRNA splicing dynamics. We use VeloVAE to infer RNA velocity from stereo-seq data. VeloVAE is a deep generative model that recovers the temporal order of cells and gene splicing dynamics in terms of the transcription, mRNA splicing and degradation rates using a pair of neural networks. The model consists of an encoder, which performs variational inference of latent cell time and state, and a decoder, which predicts unspliced and spliced counts using an ODE model.

All datasets were preprocessed using the scVelo preprocessing pipeline. Specifically, we selected 2000 top highly variable genes while keeping all genes involved in a predetermined list of cell-cell interactions. Then, we trained VeloVAE using default hyper parameters except for the latent dimension and batch size. We used a latent dimension of 5 for stereo-seq E9.5 and 10 for all other datasets. As for batch size, we set it to default (128) for stereo-seq 9.5, increased it to 256 for stereo-seq E12.5, further increased it to 512 for stereo-seq E10.5, E11.5, E13.5, E14.5, E15.5, and set it to 2048 for stereo-seq E16.5. The batch size is generally proportional to the cell number. After training, we extract the inferred parameters and compute RNA velocity using the analytical form: *ν_u_* = *α* − *β_u_*, *ν_s_* = *βu* − *γs*, using predicted u and s values.

Next, we feed RNA velocity from VeloVAE to VeloCytoSignal to compute the LR velocity. Since gene expression is the sum of spliced and unspliced counts, we can rewrite the CytoSIgnal test statistic as follows:

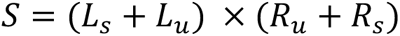

Since the statistic *L* and *R* are smoothed based on spatial neighbors, we also apply a spatial smoothing on RNA velocity estimates from VeloVAE following the exact strategy. Next, it’s straightforward to calculate the time derivative of *S* at each location following the product rule of differential calculus:

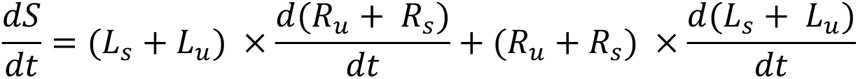

After calculating LRvelo estimates at each location, we take the average across each location’s DT neighbors to get LRvelo estimates at each spatial neighborhood.

### Visualization of signal sender-receiver cell pairs and dynamics of signaling in 3D

We adopt a public R package, *plot3D*, as a base dependency for making 3D visualizations. For making the 3D edge plot, we first export all significant receiver-sender pairs identified for the ligand-receptor interactions of interests. In the 3D space, we then create two vertically aligned scatter plots of cell locations for showing the significant receiver and sender cells respectively. To highlight the cells involved in the interaction, we set the transparency of sender cells and receiver cells higher than other cells in both scatter plots separately. Lastly, for each receiver-sender pair, we connect them by a straight line. For a cleaner visualization, we randomly downsample the cell pairs in practice.

For creating the 3D velocity plot of an interaction of interest, we first bin the cell locations into a desired number of hexagons. We then take the average of the LRvelo estimates of the cell within each hexagon. Next, we plot all cells using their spatial locations. Then we draw an arrow that vertically points from the center of each hexagon to the z-axis coordinate at the average representation of velocity. This way, the positive regional velocity results in an arrow pointing upwards and the negative velocity points downwards. Meanwhile, we color positive arrows red and negative arrows blue, to make it easier to discern the sign of the signaling velocity.

Additionally, we draw a light gray dot for regional velocity that happens to be equal to zero. For a better visualization experience, we elevate all arrows with an arbitrary height so that they don’t overlap with the scatter plot layer.

### Benchmarking analyses with previous methods

For quantifying whether the results of CytoSignal and previous methods align with PLA results, we adopt the concept of signal-sending and receiving cluster pairs from earlier methods and define a connectivity ratio (**r**) as the ratio between the number of significant edges (**s**) and the number of all possible edges (**t**) between a signal-sending and a signal-receiving cluster. An edge represents the interaction between two nearby cells that are either directly connected or within 200 μm of each other. Significant edges (**s**) denote the edges that end in locations within the receiving cluster that are identified as statistically significant by spatial permutation.

We then compared the physical distances between signal-sending and receiving cells identified by CytoSignal and previous methods. This analysis included all 332 interactions in the intersection of CellChatDB and CellphoneDB V2 and was applied to one whole mouse embryo dataset (E12.5 E1S2, 46,982 cells and 20 clusters). We ran four workflows with default parameters: CytoSignal, CellChat 2.1.0, CellChat spatial, and CellphoneDB 2.1.7. Since CytoSignal directly outputs signal-receiving cells and their corresponding signal-sending cells, we directly calculated their physical distances. Previous methods present the p-values of each interaction for each signal-sending and receiving cluster pair. To obtain the physical distances between each signal-sending and receiving cell pair from cluster-level inferences, we randomly sampled 1000 cells from the sending and receiving cluster individually and matched them into 1000 pairs. This was done for each significant interaction identification and each cluster pair. We calculated the spatial distance between each cell pair and finally formed the distribution.

We benchmarked the time and memory usage of CytoSignal and previous methods using their default settings. Each method was benchmarked using the same computational resources on the Fedora Linux OS platform with a single Intel Xeon 3.00GHz core and 32 GB of memory. Due to the difference in the database content of the methods we benchmark, we selected a total of 50 interactions that are in the intersection of CellChatDB and CellPhoneDB V2. For the 50 entries, we randomly selected 16 contact-dependent interactions, 17 diffusion-dependent interactions with single-molecule receptors, and 17 diffusion-dependent interactions with complex receptors. For CytoSignal, we benchmarked the imputation of ligands and receptors, the calculation of LRScores, and the permutation test as the main workflow. For CellChat 2.1.0, we benchmarked the computation of communication probabilities and the aggregation of the network. The same applies to the CellChat workflow with spatial information provided. These three sets of tests were performed in R 4.3.1, where we captured the time usage with package *microbenchmark* 1.4.10 and the RAM usage with package *peakRAM* 1.0.2. CellPhoneDB 2.1.7 is presented as a command-line interface (CLI) tool, thus the time usage was captured with shell command “time” whereas the memory usage report was generated with Python package *cmdbench* 0.1.13.

### Mouse experiments

C57BL/6J (JAX000664) mice were acquired from the Jackson Laboratory. All mice were housed in the animal facility accredited by the Association for Assessment and Accreditation of Laboratory Animal Care (AAALAC), located in the Behavioral and Biological Sciences Building (BBSB) of the University of Texas Health Science Center at Houston. Animal rooms were climate controlled to provide temperatures of 21+/-1°C, 30-65% of humidity on a 12 hr, light/dark cycle (lights on at 0600 Central Standard Time). Mice were housed in individually ventilated cages (Tecniplast, Buguggiate, Italy) and in a specific pathogen-free condition. We have complied with the ARRIVE 2.0 guidelines. Access to water and food (irradiated LabDiet 5053, St. Louis, Missouri) was ad libitum. All procedures complied with the Guide for the Care and Use of Laboratory Animals and approved by the University of Texas Health Science Center at Houston’s Animal Welfare Committee (AWC), protocol AWC-21-0070. For prenatal experiments, male C57BL/6 mice were mated to female C57BL/6 mice and the vaginal plug was checked in the morning and determined as 0.5dpf. Pregnant mice were sacrificed at embryonic day (E)12.5. Pups and fetuses were used for analysis regardless of the sex. A total of 2 pregnant female mice and 12 fetuses were used for the experiments. Mice were euthanized by over-dosage of inhalation anesthesia in a drop jar (Fluriso, Isoflurane USP, VetOne) followed by decapitation.

### Proximity ligation assay

Proximity ligation assay (PLA) was performed according to the manufacturer’s instructions (DUO92105 Sigma Aldrich). Samples were fixed in 4% paraformaldehyde overnight at 4 °C, then were cryoprotected in 30% sucrose/PBS solutions and in 30% sucrose/PBS: OCT (1:1) solutions, each overnight at 4 °C. Samples were embedded in an OCT compound (Tissue-Tek, Sakura) and cryosectioned at 14 μm using a cryostat (Leica CM1860) and adhered to positively charged glass slides (Fisherbrand ColorFrost Plus). Sections were postfixed in 4% paraformaldehyde for 15 min at room temperature, treated with 0.1% TritonX-100 for 10 min at RT for permeabilization, blocked with Duolink® blocking solution(DUO82007) at 37 °C for 1 h, and incubated with each antibodies for FGF8(Goat polyclonal, 1:200, Invitrogen, PA5-47598) and FGFR1(Rabbit polyclonal, 1:200, Abcam, ab59194), or for OSTP(Goat polyclonal, 1:200 R&D system, AF808) and CD44(Rabbit polyclonal, 1:200, Proteintech,15675-1-AP), or for IGF2 (Goat polyclonal antibody, 1:200, R&D systems, AF792) and MPRI (Rabbit polyclonal antibody, 1:200, Abcam, ab124767), for DLL1(Rabbit polyclonal, 1:200, Abcam, ab10554) and NOTCH1(Goat polyclonal, 1:200, R&D system, AF1057) or for EFNA3(Rabbit polyclonal, 1:200, Proteintech, 12480-1-AP) and EPHA5(Goat polyclonal, 1:200, R&D system, AF3037) or each negative control (using only receptor antibodies for primary antibodies) overnight at 4℃. Subsequently, sections were incubated with Probe mix [Duolink® In Situ PLA-® Probe Anti-Rabbit MINUS(DUO92005) and Duolink® In Situ PLA-® Probe Anti-Goat PLUS(DUO92003)] in a preheated humidity chamber for 1 h at 37 °C, followed by incubation with a ligation solution for 30 min at 37 °C and an amplification solution for 100 min at 37 °C. Sections were further incubated with DAPI (4ʹ,6-diamidino-2-phenylindole, 5 μg/ml, Invitrogen D1306) to stain nuclei prior to imaging. Images were captured by an automated inverted fluorescence microscope with a structured illumination system (Zeiss Axio Observer7 with ApoTome 3 system) and Zen 3.4 software. The Filter Set 112 SBP was used with excitation (Ex), beam-splitter (Bs) and emission (Em) filter wheel consisting of: Ex. 385/30, 469/38, 555/30, 631/33, 735/40, Bs. 405 + 493 + 575 + 654 + 761, Em. 425/30, 514/30, 592/25, 681/45, 788/38 nm. The objectives used were: Plan-Apochromat 10x/0.45, Plan-Apochromat 20x/0.80, and Plan-Apochromat 63x/1.40.

### Preprocessing and binning stereo-seq data

Raw data files (*FASTQ format*) sequenced with Stereo-seq for mid and late-gestation mouse embryos were downloaded from CNGBdb (https://db.cngb.org/search/project/CNP0001543/). Apart from the sequencing data, a barcode-to-position mapping file was also obtained from the Mouse Organogenesis Spatiotemporal Transcriptomic Atlas (*MOSTA :* https://db.cngb.org/stomics/mosta/download/) for each of the embryonic sections. The sequencing data was then processed using the Stereo-seq Analysis Workflow - SAW_v5.1.3. A STAR genome index was built using the GRCm38 assembly. A transcriptome reference file (*Mus_musculus.GRCm38.100.gtf*) was downloaded from Ensembl and provided as an input to the SAW pipeline along with the STAR genome index and the barcode-to-position file for Stereo-seq reads alignment.

A position-sorted and deduplicated Binary Alignment Map (BAM) file was obtained as a result of the SAW workflow for each of the Stereo-seq samples. The BAM file was used to quantify the spliced and unspliced counts using a custom Perl script provided by BGI which utilizes the BAM read tags to extract the spatial coordinates and the exonic/intronic counts for each spatial position. To get count matrices at cellular level, we used custom python codes to bin the data using the same bin size as the original analysis.

## Supplementary Figures

**Supplementary Fig. 1.**
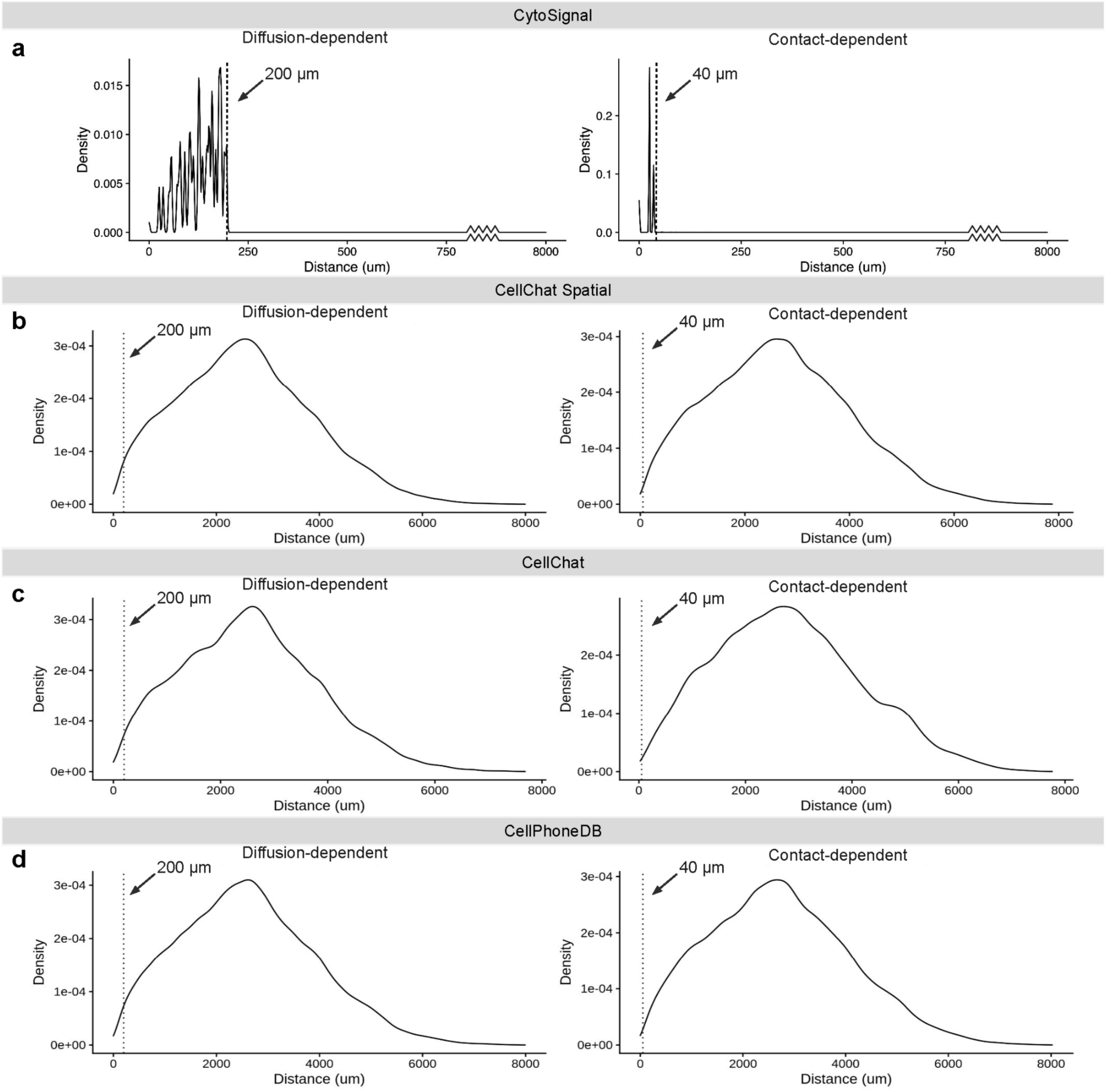
Benchmarking the physical distances between significant signal-sending and signal-receiving locations inferred by CytoSignal and previous methods. **a**, Probability density of significant signaling location pairs as a function of physical distance identified by CytoSignal. The x-axis is the physical distance in µm and the y-axis is the density of pairs of signal-sending and signal-receiving locations counts at each distance. Note that the x-axis is truncated between 750 µm to 8000 µm because the density is 0 within this range. **b**, Probability density of signaling location pairs identified by CellChat Spatial as a function of their physical distances. **c**, Probability density of signaling location pairs identified by CellChat as a function of their physical distances. **d**, Probability density of signaling location pairs identified by CellPhoneDB as a function of their physical distances.

**Supplementary Fig. 2.**
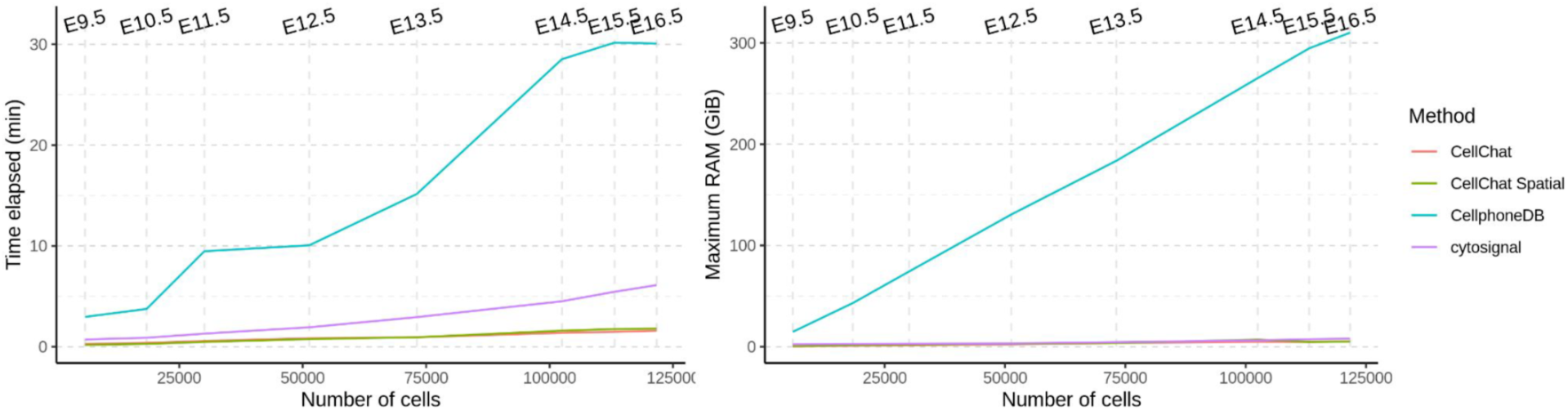
Benchmarking time and memory usage of CytoSignal and previous methods. Each method is tested on a total of 8 stereo-seq datasets. The x-axis is the number of cells in each dataset. The left panel’s y-axis is the time used and the right panel’s y-axis is the memory used. Each method is shown as a different line.

**Supplementary Fig. 3.**
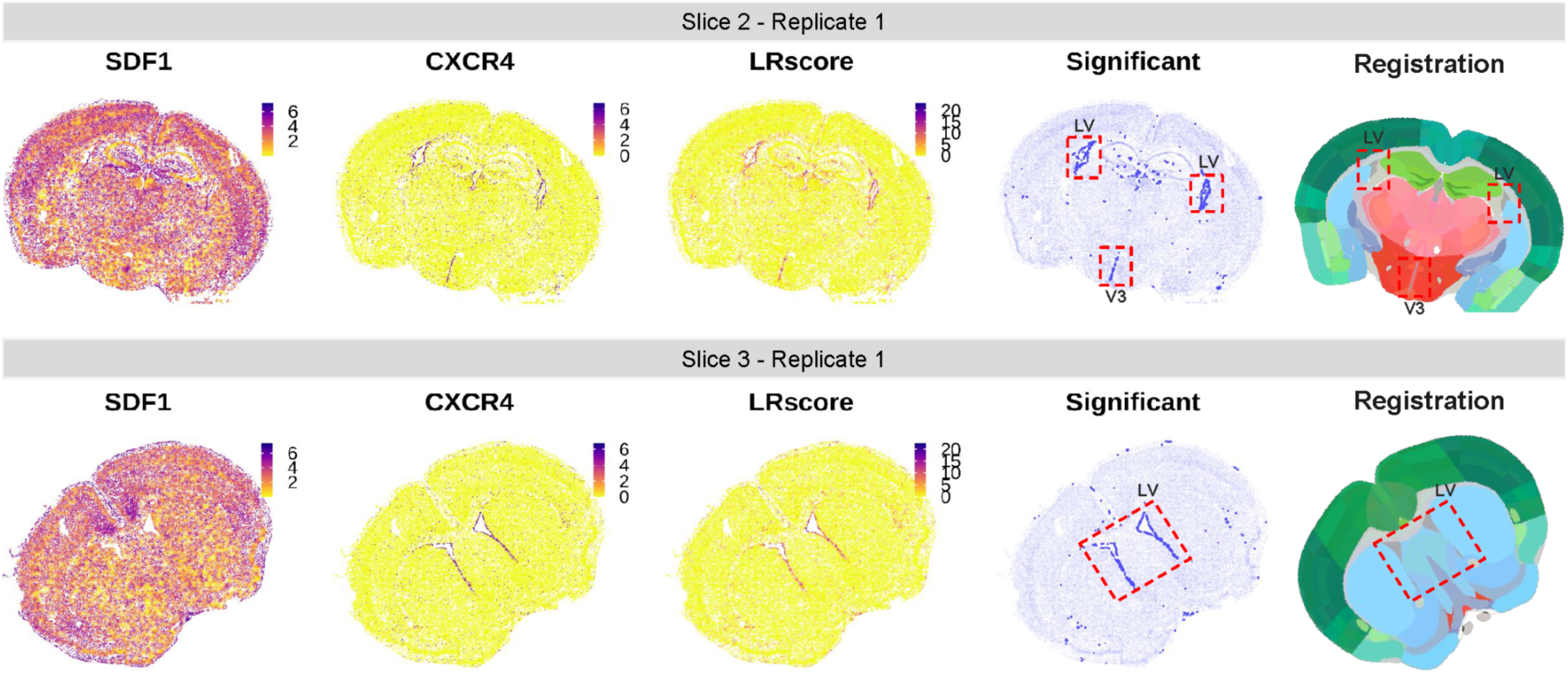
CytoSignal identifies significant interactions closely related to adult neurogenesis in MERFISH data. From left to right: Ligand (Sdf1) and receptor (Cxcr4) expression data, LRscore, inferred locations with significant signaling activity, and brain region annotations for each location. LV: lateral ventricle, V3: third ventricle.

**Supplementary Fig. 4.**
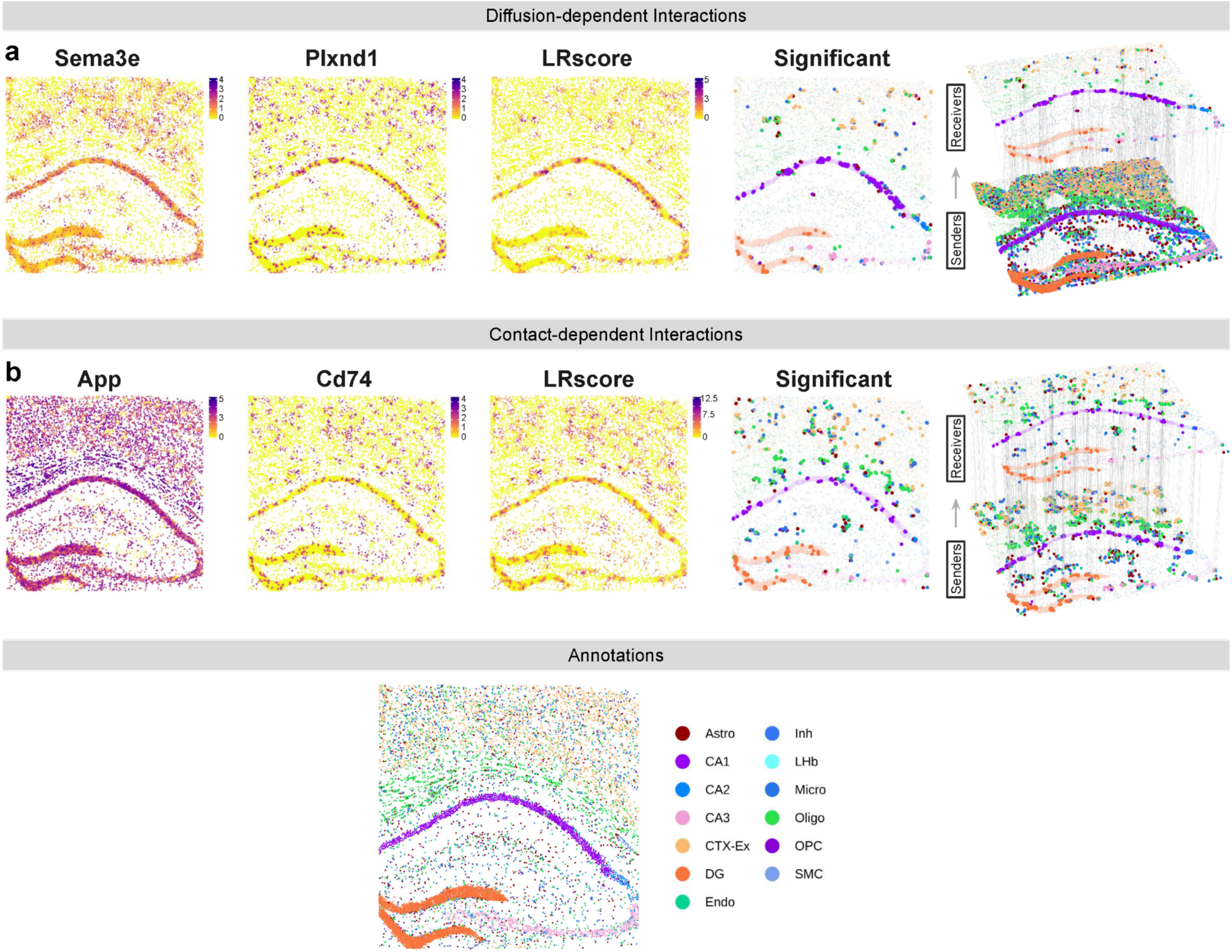
CytoSignal identifies signaling interactions related to amyloid from STARmap PLUS data. **a,** Ligand (Sema3e) and receptor (Plxnd1) expression data, LRscores, inferred locations with significant signaling activity, and inferred signal-sending and signal-receiving cells. **b,** Ligand (App) and receptor (Cd74) expression data, LRscores, inferred locations with significant signaling activity, and inferred signal-sending and signal-receiving cells.

